# Fluorescence-Based Detection of Fusion State on a Cryo-EM Grid using Correlated Cryo-Fluorescence and Cryo-Electron Microscopy

**DOI:** 10.1101/388579

**Authors:** Lauren Ann Metskas, John A. G. Briggs

**Affiliations:** Structural and Computational Biology Unit, European Molecular Biology Laboratory, Heidelberg, Germany; Structural Studies Division, MRC Laboratory of Molecular Biology, Cambridge, United Kingdom

## Abstract

Correlated light and electron microscopy (CLEM) has become a popular technique for combining the protein-specific labeling of fluorescence with electron microscopy, both at room and cryogenic temperatures. Fluorescence applications at cryo-temperature have typically been limited to localization of tagged protein oligomers due to known issues of extended triplet state duration, spectral shifts, and reduced photon capture through cryo-CLEM objectives. Here, we consider fluorophore characteristics and behaviors that could enable more extended applications. We describe how dialkylcarbocanine DiD and its autoquenching by resonant energy transfer can be used to distinguish the fusion state of a lipid bilayer at cryo-temperatures. By adapting an established fusion assay to work under cryo-CLEM conditions, we identified areas of fusion between influenza virus-like particles and fluorescently labeled lipid vesicles on a cryo-EM grid. This result demonstrates that cryo-CLEM can be used to localize functions in addition to tagged proteins, and that fluorescence autoquenching by resonant energy transfer can be incorporated successfully into cryo-CLEM approaches. In the case of membrane fusion applications, this method provides both an orthogonal confirmation of functional state independent of the morphological description from cryo-EM and a way to bridge room-temperature kinetic assays and the cryo-EM images.

## Introduction

Correlated light and electron microscopy (CLEM) provides its users both the specificity of fluorescence microscopy and the resolution of electron microscopy (EM), and has now become an established technique with many variations (1). Room-temperature applications permit better fluorescence optics but limit the EM to resin-embedded samples; conversely, cryo-temperature applications allow the full power of cryo-EM but at the expense of the numerical aperture of the fluorescence objective (2). Despite its optical drawbacks, interest and applications for cryo-fluorescence microscopy (cryo-FM) are growing due to the potential to correlate with cryo-EM; the advantages of cryo-freezing as a means of rapid, chemical-free sample fixation; and the possibility of reduced photobleaching at low temperature (3–5).

One area where cryo-fluorescence is still lacking is the expansion to applications that draw on unique dye photophysics such as Förster Resonance Energy Transfer (FRET) and stochastic photobleaching. Such approaches are rarely used at cryo-temperatures due to optical limitations and potential temperature dependence in the behavior of the dyes. This has functionally limited cryo-CLEM to being a method for targeting a fluorophore-tagged molecule for cryo-EM imaging.

During a reaction or process in which no protein is added or lost, localization-based methods will not give a distinguishable signal change in cryo-CLEM. Such processes include lipid membrane events like fusion and fission. While in a process like endocytosis individual proteins can be tagged to organize images in time (6), fluorescence localization cannot determine whether two membranes in tight apposition are exchanging lipids or not, or exactly when a hemifusion stalk becomes a fusion pore. In such cases a chemical reporter is necessary to define the point at which reactions begin (7–9).

In this study, we have adapted a standard fusion assay to cryo-CLEM, employing an application of autoquenching based on resonant energy transfer (RET). By doing so, we apply cryo-CLEM to localize a functional event rather than a protein, drawing upon the expanded fluorophore uses that have been mostly neglected in cryo-CLEM applications. We use the model system of influenza virus fusion, which has been well-studied separately by fluorescence kinetics (8) and cryo-EM and tomography (10, 11). The use of the cryo-fluorescence signal to localize a function allows distinction between apposition and hemifusion, allowing targeted imaging of hemifusion and permitting more accurate assignment of micrographs into their appropriate functional state.

## Fluorophore selection criteria and controls

Assays that monitor membrane fusion are usually based on tracking lipid mixing, content release, or both. In lipid mixing assays, a fluorophore is typically embedded in one membrane; upon fusion, the fluorophore is diluted in the expanded membrane and gives a fluorescence signal, typically dequenching. Quenching can occur by multiple mechanisms, including collision (dynamic quenching), formation of a dark complex (static quenching), or resonant energy transfer (RET), and can be a product of contact with the fluorophore itself (autoquenching) or with other small molecules (12). While room-temperature, solution-based applications can draw from a wide variety of molecules and mechanisms, cryo-FM places specific demands on fluorophores and their photophysical behavior.

There are many considerations for choosing a fluorophore for cryo-CLEM applications. At cryo-temperatures (−180 to −135 C), macromolecular diffusion will not occur. The cryo-EM grid has an unprotected surface that can nonspecifically bind or activate fluorophores, increasing background signal. Cryo-temperatures have strong effects on fluorophores: excitation and emission spectra can shift or change in width (13, 14); fluorophores can spend large portions of time trapped in the triplet state (2, 15); and some photo-switchable fluorophores fail to work as expected or experience decreased quantum yields (16, 17). Finally, due to the optical and experimental limitations of cryo-FM, the labeled sample is visualized with a low-numerical aperture objective and a low enough illumination intensity to avoid devitrification.

Given these constraints, the fluorophores with the best potential for the study of membrane fusion by cryo-CLEM would autoquench, and could be embedded in either target or fusogen-containing membrane. The signal must be bright, and the change upon membrane fusion should be strong enough to be visible without extensive processing (a near-binary dark-bright transition). Based on these criteria, we selected two dyes for testing: octadecyl rhodamine B (R18) and the dialkylcarbocanine DiD. Fluorophores were loaded into a 1 mg/mL preparation of extruded POPC vesicles at concentrations below those required for full quenching, plunge-frozen onto EM grids, and imaged in a Leica cryoCLEM system (see SI for extended methods). R18 was tested in 100 nm POPC vesicles at a 4% concentration. In these grids, the R18 appears to have partitioned onto the carbon support layer of the EM grid and entered a high-fluorescence state at that location, leading to an unsuitably high background (Figure 1A, top). In contrast to the R18, cryo-FM images of 200 nm POPC vesicles containing 1% DiD show fluorescent punctae scattered over the carbon and holes (Figure 1A, bottom).

**Figure 1.**
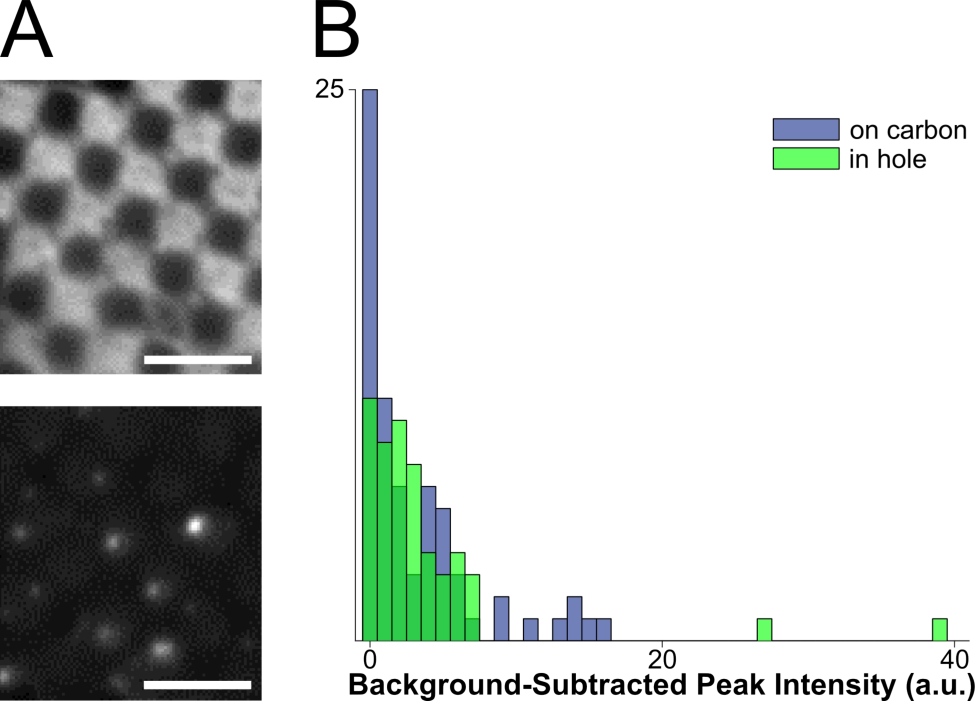
Assessment of fluorophores for labelling vesicles on cryo-EM grids. **A**: top: cryo-FM image of POPC vesicles labeled with 4% R18, showing a lawn of fluorescence intensity staining the carbon of the EM grid. Bottom: cryo-FM image of POPC vesicles labeled with 1% DiD, displaying punctate behavior. Scale bars = 5 µm. **B**: quantification of intensities for clearly separated vesicles labelled with 5% DiD, sorted by location in holes or on carbon.

After confirming that DiD labeling produced punctate behavior in cryo-FM, we determined the intensity change that could be expected from DiD dequenching upon membrane fusion. Room-temperature fusion assays are often baseline-corrected and normalized, allowing the use of fluorophores with modest intensity changes; however, such processing is error-prone in cryo-FM due to intensity variability by ice thickness, stray reflections, and a lack of negative and positive control areas within the imaging square for establishing baselines. For cryo-CLEM, therefore, both a dark base state and a large intensity change upon fusion are required. We measured the fluorescence emission of POPC vesicle preparations labeled with varying molar percentages of DiD. We observed complete dequenching at DiD concentrations of 5% and above. We confirmed that quenched DiD vesicles would be darker than unquenched vesicles regardless of fluorophore amount at relevant pHs (Figure S1A) and that DiD autoquenching could yield a roughly 10-fold intensity change upon fusion between a vesicle and a similarly-sized virion (Figure S1B).

We next assessed the consistency of small, quenched DiD vesicles that would be appropriate for fusion assays. A 1 mg/mL POPC/POPS suspension was labeled with 5% DiD, extruded through 50 nm pores, and plunge-frozen onto grids containing 50 nm tetraspec beads (SI). Grids were imaged in cryo-FM as above. Grid squares of interest were then imaged on an FEI Tecnai F20 electron microscope (SI). The cryo-FM and cryo-EM micrographs were correlated using Matlab-based cryo-CLEM correlation software (18) and FIJI (19, 20) with the Stack Focuser plugin. Finally, fluorescence pixel intensities were quantified for each lipid vesicle within the squares of interest (Figure 1B, SI methods).

These cryo-CLEM results confirmed that the fluorescence intensity of the quenched DiD liposomes is consistently low, and is not affected by adhesion to the carbon grid surface (Figure 1B). No intensity corrections were made for vesicle size variation or multiple lamellae, neither of which can be assessed prior to using the cryo-FM image for targeting EM imaging. In this context, the narrow intensity distribution in Figure 1B shows that variation in the total amount of fluorophore present in a spot does not result in strong signal heterogeneity in the quenched state, consistent with full quenching occurring at the same fluorophore concentration as at room-temperature (Figure S1).

## Localizing membrane hemifusion events in cryo-CLEM

We elected to use influenza virus-like particles (VLPs) to induce fusion; the influenza fusion mechanism is well-studied and VLP samples are well-suited for the concentration and thickness constraints of EM. VLPs were prepared by transfecting HEK 293T cells with plasmids containing the influenza A proteins hemagglutinin, M1, M2, and neuraminidase as in (21) (SI). VLPs were harvested by centrifugation through a sucrose cushion (SI), after which their mixed morphology (filaments and spheres) was maintained, and assessed for fusogenicity (SI, Figure S2). 100 nm target vesicles were prepared by extruding a 1 mg/mL suspension of 44.7% POPC, 12.3% POPE, 33.2% cholesterol, 4.8% total ganglioside, and 5% DiD in HEPES-citrate buffer (membrane composition adapted from (10)).

We prepared EM grids to assess fusion localization by cryo-CLEM. VLPs and quenched-DiD vesicles were incubated on ice at relative concentrations of 0.16 mg/mL accessible viral protein and 0.12 mM target membrane lipid to allow VLP-vesicle adhesion. The tube contents were then acidified at pH 5.1 for 60 s at room temperature, followed by addition of 10 nm gold fiducial markers and plunge-freezing in liquid ethane (SI). The total time between acidification and freezing was roughly 2.5 minutes, at which time the hemifusion reaches a plateau at room temperature (SI, Figure S2).

The EM grids were then subjected to cryo-FM, following the same workflow as with the control DiD vesicle sample above (Figure 1B). Cryo-FM imaging was performed using a ten-fold lower exposure time appropriate for the high-intensity, dequenched punctae; we therefore expected a very low signal from the quenched vesicles. Brightfield and fluorescence maps of the EM grid were correlated on-the-fly with EM micrographs collected on an FEI Tecnai T12 microscope using established SerialEM-based correlation protocols (22, 23).

Selected grid squares were imaged at a magnification at which VLPs and vesicles could be identified (3500-6000 X), allowing later evaluation of the full grid square by correlating the lower-resolution cryo-FM and cryo-EM images in post-processing (18). Because we expected fusion to dequench the DiD and yield bright cryo-FM signals, we targeted bright- and medium-intensity punctae for higher-magnification EM imaging using on-the-fly correlation at the cryo-EM (Figure 2A-B) (23). This combination of approaches allows targeted high-resolution acquisitions at spots of interest as well as lower-resolution evaluation of both targeted and untargeted areas during post-processing (Figure 2A-C).

**Figure 2.**
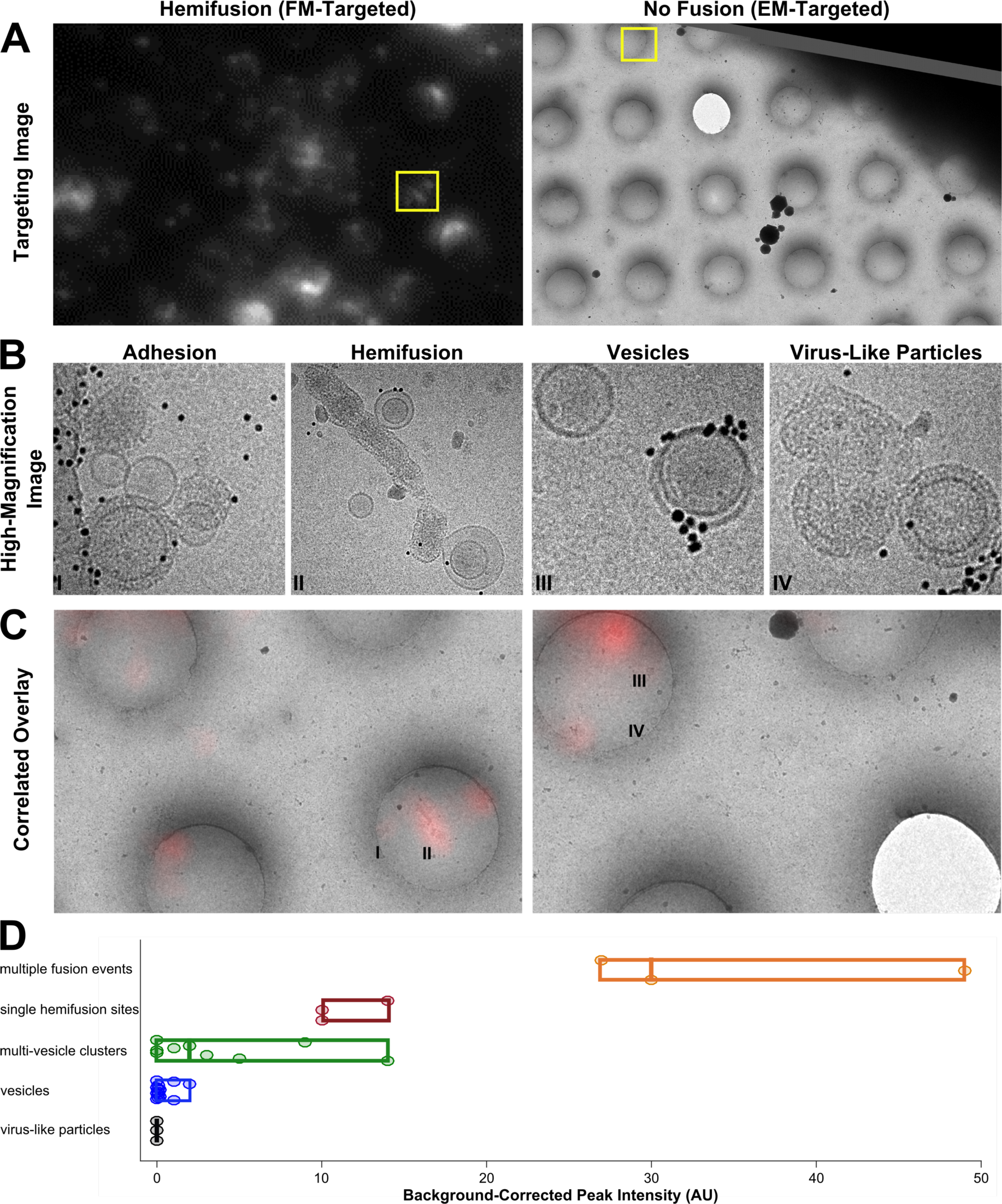
Cryo-CLEM images of different types of on-grid events and their corresponding fluorescence signals. **A**: Targeting images are used to define points of interest (yellow boxes). Left: fluorescence-based targeting of potential hemifusion sites. Right: EM-based targeting of VLPs and vesicles irrespective of fluorescence. **B**: High-magnification micrographs are taken within the targeted areas. **C**: During post-processing, fluorescence and EM maps are correlated with greater precision, and fluorescence intensities are quantified for all imaging locations. Roman numerals correspond to the high-magnification images in B. **D**: Fluorescence intensities for vesicles and virus-like particles. Events are classified visually. Circular points are single events, boxes display ranges per category, and mid-lines represent the median value for that category.

After examining the overlay of fluorescence signal with the electron micrographs, we observed that the brightest spots on the grid corresponded to aggregates or clusters of VLPs and vesicles with multiple fusion sites (Figure 2D). The intermediate intensity spots typically contained either hemifusion events or clusters of many adhesion events (Figure 2). Importantly, none of the intermediate or bright fluorescence events correlated to unfused VLPs or solitary vesicles, indicating that the DiD remains quenched in the unfused vesicles and that there is no transfer of dye outside of active membrane fusion (Figure 2B-D). Unfused VLPs and solitary vesicles gave very low fluorescent signals (Figure 2D). Considering all trends together, if all bright and intermediate fluorescence events were targeted, all hemifusion sites should be contained in the EM dataset.

## Discussion and Future Prospects

In this study, we demonstrated that the established DiD-based fusion assay (11, 24) can be incorporated into a cryo-CLEM workflow to specifically locate, target and image membrane fusion sites. By using the same fluorescence-based fusion signal as room-temperature kinetic assays (Figure S2), this method also provides a way to link the EM-based ultrastructural imaging with orthogonal kinetic and functional information. Furthermore, our use of DiD, which employs an RET-based autoquenching mechanism (25), confirms that RET-based fluorescence applications can function at cryo-temperatures under favorable conditions. Of these applications, autoquenching is particularly well-suited for cryo-CLEM because the use of a single fluorophore decreases the likelihood of spectral shifts strongly affecting the overlap integral for RET. RET applications with many dye molecules present, such as membrane fusion, are also optimal: the number of fluorophores is likely sufficient to allow the random orientation of the fluorophore dipoles to approximate the tumbling seen at room temperature, and fluorescent emission will be strong despite cryo-temperature limitations such as extended triplet state occupation, gentle illumination intensity or low numerical aperture.

One natural extension of this method will be to load DiD into viruses to locate membrane fusion within eukaryotic endosomes, as has been done at room temperature (24). This would facilitate the localization and imaging of rare fusion events within vitrified cellular samples. For the study of membrane fusion *in vitro*, our cryo-CLEM method also provides an orthogonal signal for lipid mixing, which cannot always be visually distinguished from membrane apposition in cryo-EM images. The use of fluorescence dequenching to study influenza fusion allows correlation between an image and previous kinetics studies based on similar fluorescence signals (8), ensuring that the definition of hemifusion is consistent between the kinetic and structural methods.

In conclusion, we have developed a method for localizing a specific function on a cryo-EM grid, using an application of RET in cryo-CLEM. This development allows localization of a fusion function rather than a protein of interest in cryo-FM, which can be used either to target this site for cryo-EM imaging, or to provide an orthogonal evaluation of the fusion state.

## Author Contributions

L.A.M. performed the research and data analysis. L.A.M. and J.A.G.B. designed research, interpreted results and wrote the manuscript.

## Acknowledgements

This research was technically supported by the Electron Microscopy Core Facility and the Protein Expression and Purification Core Facility at the European Molecular Biology Laboratory (EMBL, Heidelberg), and the EM facility at the MRC Laboratory of Molecular Biology. We thank M. Schorb, W. Kukulski E. Lemke, G. Paci, O. Avinoam, and Y. Bykov for helpful discussions and technical assistance with the Leica cryo-CLEM system, and M. Clarke for assistance with VLP preparations. This project was funded by the European Research Council (ERC) under the European Union’s Horizon 2020 research and innovation programme (ERC-2014-CoG 648432 – MEMBRANEFUSION) to JAGB. JAGB is an inventor on filed patents that have been licensed by Leica Microsystems and commercialized in the Leica cryo-CLEM system used in this study.

